# Intestinal epithelial adaptations to vertical sleeve gastrectomy defined at single-cell resolution

**DOI:** 10.1101/2023.05.31.543143

**Authors:** Kieran Koch-Laskowski, Ki-Suk Kim, Maigen Bethea, Kelly N. Z. Fuller, Darleen A. Sandoval, Praveen Sethupathy

**Author notes:** Indicates co-senior authorship.

## Abstract

The gut plays a key role in regulating metabolic health. Dietary factors disrupt intestinal physiology and contribute to obesity and diabetes, whereas bariatric procedures such as vertical sleeve gastrectomy (VSG) cause gut adaptations that induce robust metabolic improvements. However, our understanding of these adaptations at the cellular and molecular levels remains limited. In a validated murine model, we leverage single-cell transcriptomics to determine how VSG impacts different cell lineages of the small intestinal epithelium. We define cell type-specific genes and pathways that VSG rescues from high-fat diet perturbation and characterize additional rescue-independent changes brought about by VSG. We show that Paneth cells have increased expression of the gut peptide Reg3g after VSG. We also find that VSG restores pathways pertaining to mitochondrial respiration and cellular metabolism, especially within crypt-based cells. Overall, our study provides unprecedented molecular resolution of VSG’s therapeutic effects on the gut epithelium.

## Introduction

Energy homeostasis is maintained by physiological activity coordinated across organ systems. A significant contributor is the gastrointestinal tract, which serves as the primary site for nutrient intake amidst a dynamic mechanical, chemical, and microbial environment. Systemic energy balance relies in part on the gut’s ability to adapt to ever-changing conditions, an endeavor initiated by the intestinal epithelium.

Gut epithelial cells serve as a critical interface with the luminal environment. Lining the small intestine, they form crypt and villus structures that continuously renew from proliferating stem cells at the crypt base. From there, unique lineages differentiate and carry out specialized functions throughout the crypt-villus axis. Major cell types include: enterocytes, which comprise the majority of the small intestinal epithelium and facilitate nutrient digestion and absorption; goblet and tuft cells, which secrete mucus and contribute to mucosal immunity, respectively; Paneth cells, which support the crypt niche and generate antimicrobial peptides; and lastly enteroendocrine cells (EECs), which release a variety of hormones in response to luminal stimuli to help coordinate whole-body metabolism^1, 2^. Given the vast heterogeneity both between and within these lineages, a growing number of studies have leveraged single-cell transcriptomic technology to deepen our understanding of the mechanisms underlying gut health and metabolic homeostasis^3, 4^.

Perturbations in the intestinal epithelium have been linked to the pathogenesis of metabolic disease. For example, recent single-cell investigations have reported early and advanced epithelial maladaptations following consumption of diets high in fat and/or sugar^5, 6^. These studies are part of a larger effort to identify novel therapeutic targets for diet-induced obesity, a complex phenotype associated with serious comorbidities, such as type 2 diabetes; cardiovascular, liver, and musculoskeletal diseases; and mental health disorders^7^. The growing prevalence of obesity and its collective impact on global mortality impose disproportionate burdens on different facets of society^8^. Thus, further research is urgently needed to determine how the gut may be adapted to improve overall public health.

Among the current treatment strategies for obesity and metabolic disease, bariatric surgery produces the most profound effects on weight loss and other metabolic parameters, highlighting the therapeutic potential of the gut^9–11^. One of the most commonly performed bariatric procedures is vertical sleeve gastrectomy (VSG), which achieves significant metabolic improvements within one year post-surgery^12^. Despite its effectiveness, VSG is an invasive procedure not without risk, involving resection of ∼80% of the stomach along the greater curvature. This anatomical manipulation is known to induce small intestinal epithelial adaptations, particularly within the EEC lineage, that have been previously investigated by us^13^ and others^11^ for their possible roles in ameliorating metabolic disease. However, a comprehensive picture of how other cell types of the small intestinal epithelium respond to VSG has yet to be established.

Here, we aimed to define the overall cellular and molecular landscape of the small intestinal epithelium upon treatment of diet-induced obesity by bariatric surgery. Using a single-cell transcriptomic approach with a validated murine model of VSG, we identified all major cell lineages along the crypt-villus axis and noted cell type-specific genes and pathways rescued by VSG following dietary perturbation. Our results provide greater resolution in localizing changes previously observed after VSG (such as recovered expression of the gut peptide Reg3g^14^). We also reveal nuclear and mitochondrial genes involved in cellular respiration that are rescued in crypt-based lineages. Altogether, this unprecedented view highlights how adaptations among specific cell types may affect gut epithelial homeostasis, taking one step closer towards the discovery of more targeted, less invasive treatment strategies for metabolic disease.

## Results

### VSG produces robust metabolic improvements in a validated mouse model

To better understand how diet and bariatric surgery impact the small intestinal epithelium, we first initiated male C57BL/6J mice on a 16-week high-fat diet (HFD) regimen followed by surgical intervention (**Figure 1A**). Upon induction of diet-induced obesity, HFD fed mice underwent vertical sleeve gastrectomy (VSG), resulting in improved metabolic outcomes compared to their sham-operated counterparts as expected from previous studies in both murine models^15, 16^ and human patients^10, 17^. Within one-week post-surgery, HFD VSG animals lost significantly more body weight than HFD sham mice, reaching weights that approached those of a parallel group of sham mice fed a low-fat diet (LFD) (**Figure 1B**). Both HFD groups consistently outweighed the LFD mice in fat and lean mass, while weight loss induced by VSG was attributable to reduced fat but not lean mass (**Figure 1C**). As seen previously^15, 18, 19^, transient differences in food intake occurred following surgery but ultimately waned until all animals consumed similar amounts by three weeks post-surgery; however, HFD sham mice had higher energy intake than LFD-fed mice due to differences in dietary caloric density (**Figure 1D**). While HFD sham mice demonstrated robust oral glucose intolerance relative to the LFD group, VSG animals showed improved blood glucose levels comparable to those observed in LFD fed mice (**Figure 1E**). Altogether, the metabolic improvements observed here in body weight/composition and glucose tolerance align with expectations established in the bariatric surgery field and confirm our experimental cohort as a valid model of murine VSG.

**Figure 1:**
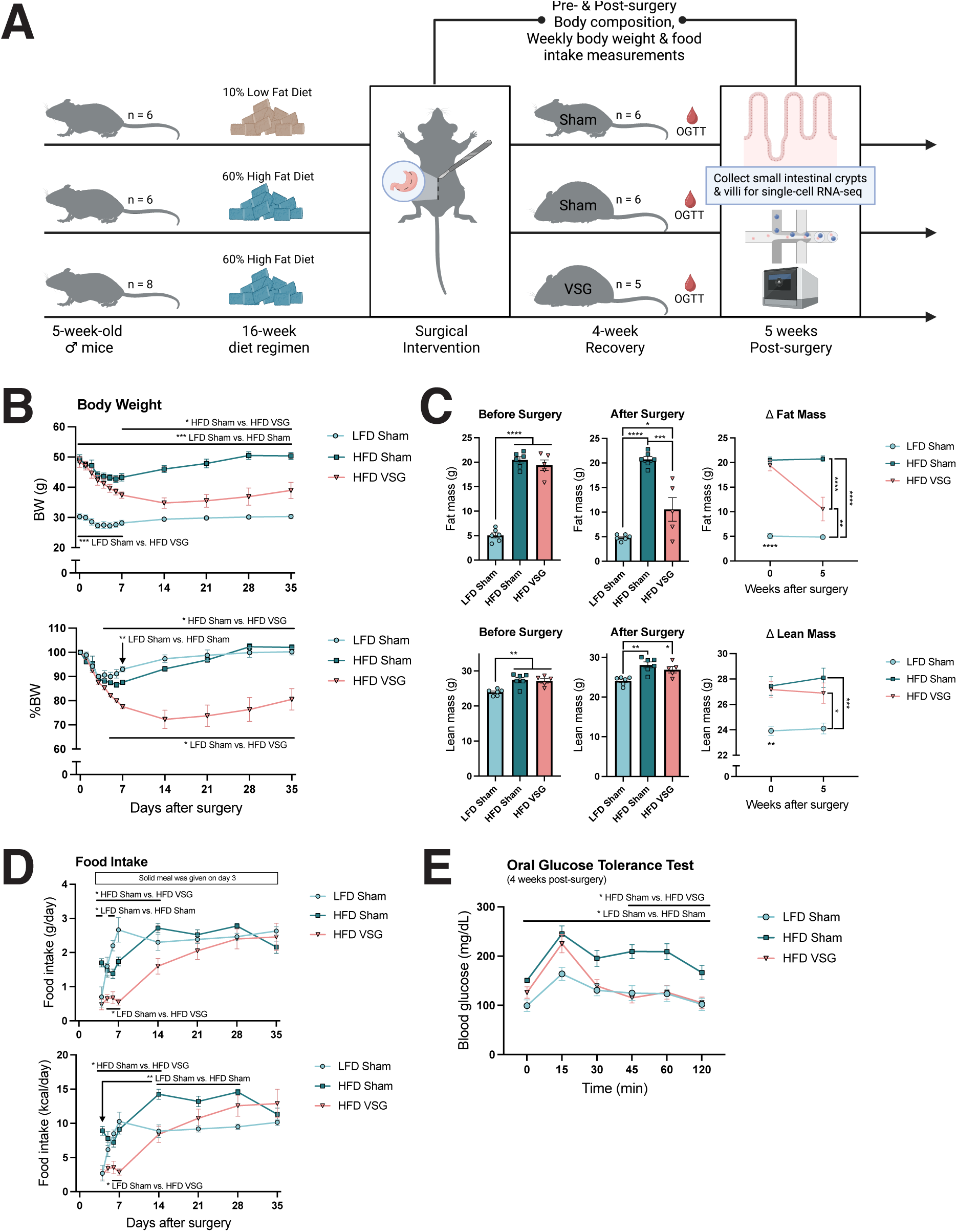
Vertical sleeve gastrectomy produces robust metabolic improvements in a validated mouse model. (A) Experimental design. Male C57BL/6J mice were maintained on 60% HFD (n=14) or 10% LFD (n=6) for 16 weeks prior to surgical intervention. HFD-fed animals underwent VSG (n=5) or a sham procedure (n=6), while LFD-fed animals were sham treated (n=6). At five weeks post-surgery, mice were overnight fasted, and small intestinal crypts and villi were subsequently collected for single-cell RNA-seq. (B) Body weight and percent body weight change. Animals were weighed daily for one week following surgery and weekly thereafter. (C) Body composition. Fat and lean mass were measured before surgery and five weeks post-surgery. (D) Food intake. Animals recovered from surgery on a liquid diet and were returned to their original solid diet at three days post-surgery. (E) Oral glucose tolerance testing. Mice were briefly fasted and tested four weeks after surgery. Data depict mean ± SEM analyzed via one- or two-way ANOVA or mixed effects analysis followed by Tukey’s post hoc testing. * *P* < 0.05, ** *P* < 0.01, *** *P* < 0.001, **** *P* < 0.0001.

### Dietary and surgical interventions induce minimal changes in intestinal epithelial morphometry and cellular composition

With substantial weight loss observed over five weeks after surgery, we next collected small intestinal tissue samples to assess how diet and surgery affect the gross morphometry and cellular composition of the epithelium. Focusing on the jejunum as a key site for nutrient absorption, we found no changes in crypt depth between HFD and LFD sham animals, consistent with a previous long-term obesogenic diet study in mice^6^. While VSG induced a slight increase in crypt depth, this trend was non-significant (**Figure 2A**). Villus height tended to rise with HFD feeding and showed an additional non-significant increase with VSG (**Figure 2B**).

**Figure 2:**
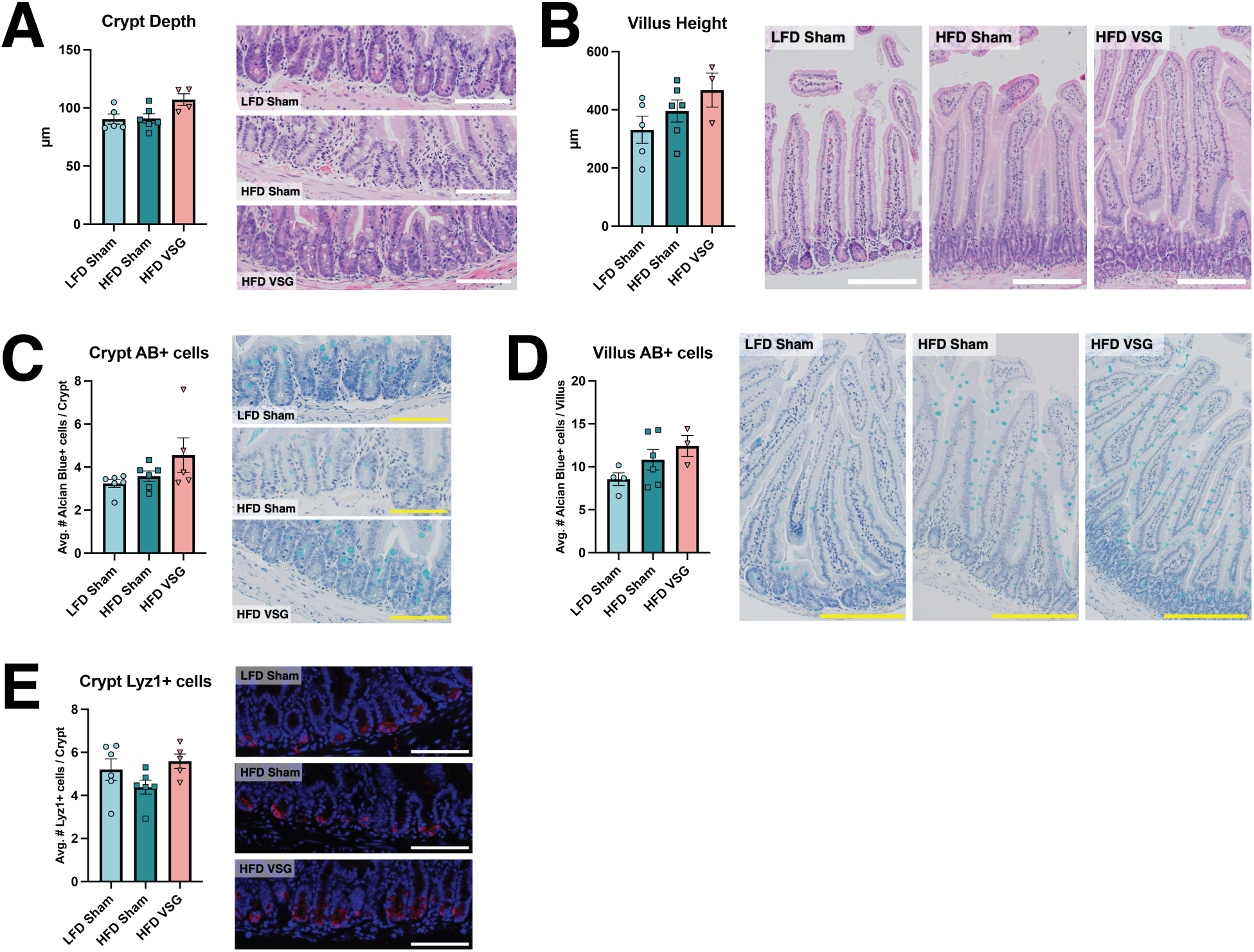
Intestinal epithelial morphometry and cellular composition show minimal changes by diet and surgery Measurements and representative histological images for. (A) Crypt depth (B) Villus height (C) Crypt Alcian blue+ cell counts (D) Villus Alcian blue+ cell counts (E) Crypt Lyz1+ cell counts Scale bar = 100µm (crypts), or 200 µm (villi). Data depict mean ± SEM, analyzed via one-way ANOVA followed by Tukey’s post hoc testing.

These findings generally replicate our recent murine VSG study^13^, in which we observed no major differences in small intestinal epithelial morphometry between VSG and sham animals. Given the emphasis we and others have already placed on surgery-induced adaptations within the enteroendocrine lineage^11, 13^, we decided to expand our focus here on how VSG affects other secretory cell types, specifically goblet and Paneth cells. Analogous to our morphometric findings, we observed modest increases in alcian blue-stained goblet cells in the crypts and villi of HFD VSG mice (**Figure 2C-D**). For crypt-based Paneth cells, we saw a mild decrease in Lyz1+ cells in HFD sham mice compared to LFD animals, which was rescued by VSG (**Figure 2E**). Overall, our histological results revealed subtle intestinal epithelial responses to different dietary and surgical contexts.

### Single-cell transcriptomic analysis defines the epithelial molecular landscape at high resolution following VSG

To delve further into how the gut adapts to a chronic HFD and treatment by VSG, we leveraged single-cell RNA-sequencing to perform, to our knowledge, the first cell type-specific investigation of the small intestinal epithelial transcriptome following bariatric surgery. Upon conclusion of our dietary and surgical interventions, we isolated jejunal epithelial samples from each of the three experimental groups (pooled from at least two biological replicates per group) and submitted separate crypt- and villus-enriched single-cell suspensions for sequencing (**Figure 1A**). The resulting datasets were quality control filtered, integrated, normalized, and analyzed through a series of bioinformatic steps (**Figure 3A**), which are detailed in the Methods section. Following computational exclusion of contaminating ambient RNA, doublets, and low-quality cells, our overall dataset comprised 24,511 cells across 19 distinct clusters. Highly enriched genes in each of these clusters were then overlapped with known markers of different intestinal epithelial lineages (as annotated in a previous single-cell survey of the small intestine^3^) to assign each cluster to a specific cell type (**Figure 3B**). Two clusters were defined as immune cells and subsequently removed to focus downstream analyses on epithelial cell types. This left a total of 21,844 intestinal epithelial cells among 17 clusters. Importantly, high-quality cells from each dietary (HFD or LFD) or surgical (VSG or sham) condition, as well as each compartment (crypts or villi), were represented proportionally throughout the dataset (**Supplementary Figure 1**). We established confidence in the assigned cell type identities by analysis of specific marker genes, including *Sis*+ enterocytes, *Lgr5*+ stem cells, *Muc2*+ goblet cells, *Lyz1*+ Paneth cells, *Dclk1*+ tuft cells, and *Chga*+ EECs (**Figure 3C**). Furthermore, the cell type assignments aligned with expectations regarding cell cycle state and differentiation status; stem and progenitor cells expressed markers of active cell cycling whereas differentiated lineages displayed greater maturation by pseudotime analysis (**Figure 3D**). With this robust single-cell dataset in hand, we proceeded to study how HFD and VSG impact gene expression across each gut epithelial cell type.

**Figure 3:**
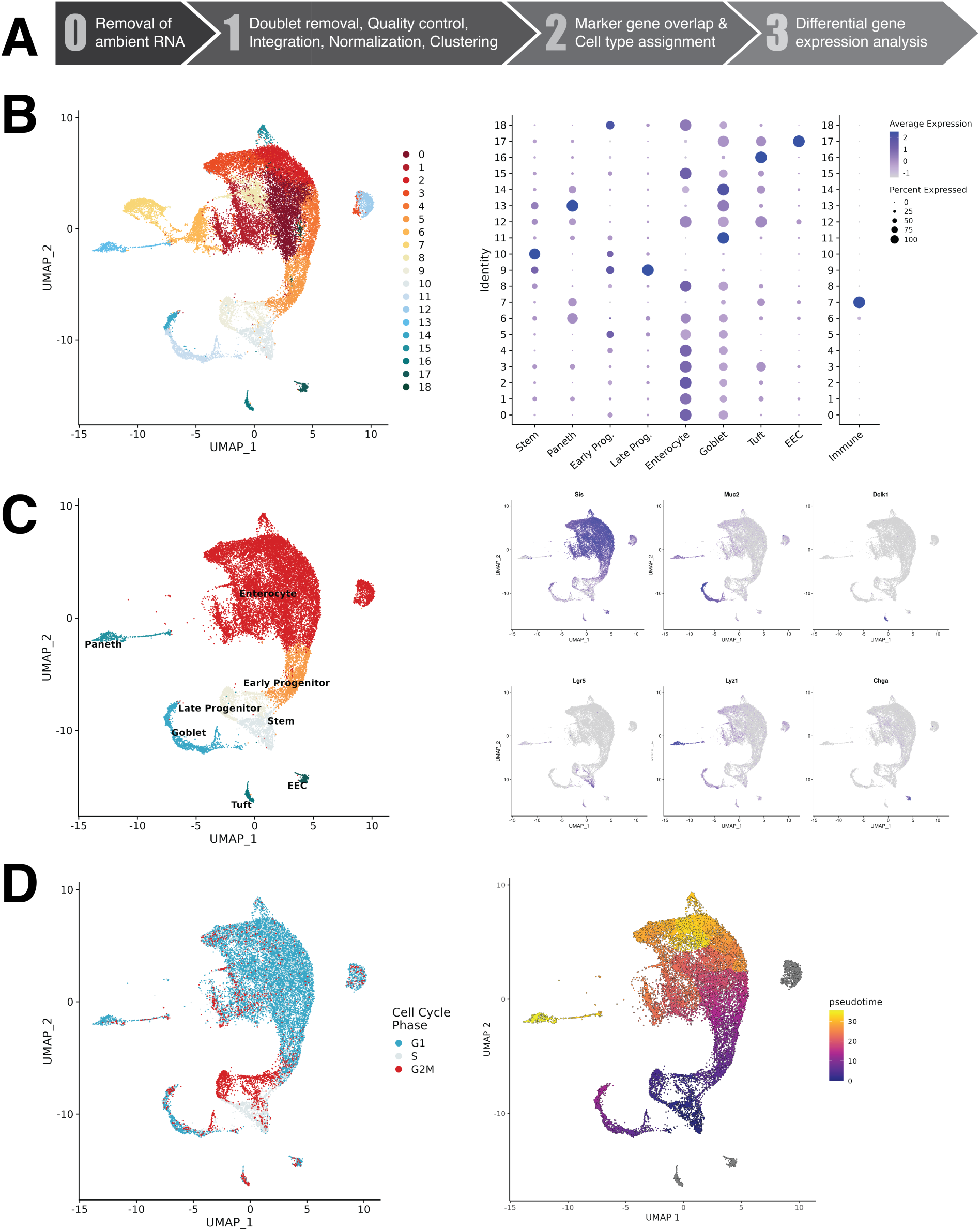
Single-cell RNA-sequencing defines the intestinal epithelial molecular landscape following bariatric surgery. (A) Schematic representation of the present single-cell RNA-seq bioinformatic workflow. (B) Uniform Manifold Approximation and Projection (UMAP) visualization of 24,511 cells across 19 distinct clusters (left), and dot plot overlap of known epithelial lineage markers with highly enriched genes from each cluster (right). (C) UMAP visualization of finalized dataset comprising 21,844 cells and their respective lineage classifications after exclusion of 2 immune cell clusters (left), and overlays of specific lineage marker gene expression (right). (D) Projections of cell cycle marker expression (left), and pseudotime trajectory predictions (right).

### Differential expression analysis highlights cell type-specific genes perturbed by HFD and rescued by VSG

For each individual cell type cluster, we asked two questions: (1) what genes are differentially expressed by HFD, and (2) what genes change in expression with VSG? This yielded two lists of differentially expressed genes (DEGs) per cluster, which we compared to see if the same genes downregulated by HFD were upregulated by VSG, and vice versa. We grouped these overlapping DEGs into one of two categories, termed “rescue” or “specificity.” “Rescue” was defined as the fraction of DEGs altered by HFD and changed in the opposite direction by VSG. “Specificity” was defined as the fraction of DEGs changed by VSG that fall into the rescue category (distinct from VSG-induced DEGs unrelated to dietary perturbation) (**Figure 4A**). To compare the extent of “rescue” and “specificity” across different cell types, we first applied filtering criteria to each DEG list (*P* < 0.05, *P*adj < 0.20, log2FC > |0.50|). Among all clusters localized to the crypt compartment, 734 genes were differentially expressed after HFD (646 down, 88 up), and 912 after VSG (772 up, 140 down). In villus-relevant clusters, there were 1496 HFD DEGs (669 down, 827 up), and 4714 VSG DEGs (2314 up, 2400 down). Visualizing these proportions in a cluster-specific manner revealed higher overall levels of rescue and specificity in crypts compared to villi, and this pattern was driven predominantly by genes downregulated by HFD and upregulated by VSG (**Figure 4B, Supplemental Figure 2**). Specific clusters that exemplified this trend include crypt-based stem and Paneth cells (as opposed to villus enterocyte clusters) (**Figure 4C**). And while the EEC lineage showed comparable degrees of rescue and specificity between crypt and villus compartments, we found that most genes rescued in EECs were distinct between crypts and villi, with the few shared being predominantly mitochondrial-encoded (**Supplemental Figure 3A-B**).

**Figure 4:**
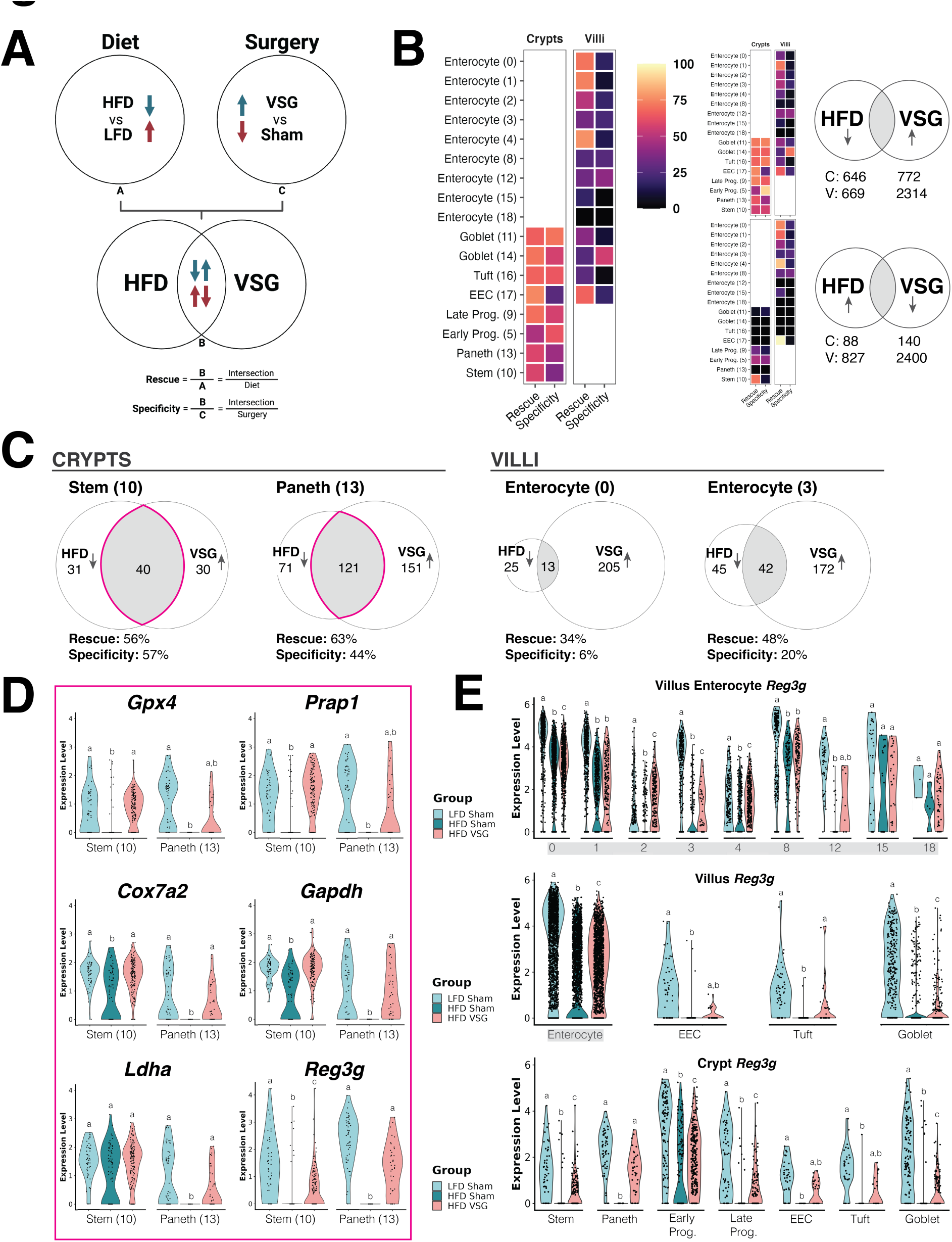
Differential expression analysis highlights genes perturbed by diet and rescued by surgery at the single-cell level. (A) Venn diagram comparison of differentially expressed genes (DEGs) between diet and surgery. Overlapping genes within the intersection are altered in opposite directions across conditions. Comparing the number of intersecting genes with the total number of DEGs within a given condition yields two fractions defined as ”Rescue” and “Specificity.” (B) Heatmap of rescue and specificity across all cell clusters, with warmer colors highlighting greater proportions. Additional plots to the right illustrate contributions from different DEG directionalities as well as the overall number of DEGs summed across the crypt or villus compartments. Detailed rescue and specificity calculations for all clusters are included in Supplementary Figure 2. (C) Venn diagrams of specific crypt- and villus-based clusters with DEGs downregulated by HFD and upregulated by VSG. (D) Violin plots of stem and Paneth cell gene expression across experimental groups. These represent a subset of DEGs rescued in the stem or Paneth clusters (or both), as indicated by the outlined intersections from (C). The full list of DEGs can be found in Supplementary Figure 3C. (E) Violin plots of *Reg3g* expression across experimental groups in all cell clusters of the crypts and villi. (D) & (E) Different letters indicate a statistically significant difference (*P* < 0.05 & *Padj* < 0.20) by MAST.

Given the critical role of the crypt in maintaining overall intestinal epithelial homeostasis^20^, we further examined the genes rescued in the stem and Paneth clusters (**Supplemental Figure 3C**). In both cell types, genes related to cellular metabolism, such as *Cox7a2* and *Gapdh*, were rescued to a similar extent (**Figure 4D**). Other genes were rescued by VSG to a greater extent in stem versus Paneth cells, such as *Gpx4*, which encodes an enzyme that mitigates harmful lipid peroxidation^21^, and *Prap1*, which codes for a protein that protects gut epithelial cells from apoptotic insults^22^. Genes that showed more Paneth-centric rescue effects included *Ldha*, which encodes a key enzyme in glycolysis leading to the production of lactate^23^, and *Reg3g*, which codes for an antimicrobial peptide that was recently shown to be required for metabolic improvements induced by VSG or a fiber-enriched diet^14^. (**Figure 4D**). We sought to leverage our single-cell data to follow up on this finding and narrow down the intestinal epithelial lineage(s) that likely drive increased *Reg3g* expression following VSG. Across nearly all clusters spanning the crypt-villus axis, *Reg3g* was downregulated by HFD; however, *Reg3g* expression returned to LFD-comparable levels most prominently in Paneth cells and villus tuft cells after VSG (**Figure 4E**).

### Pathways enriched among rescued genes underscore diet- and surgery-induced changes in nutrient absorption and metabolic function

To comprehensively survey the biological relevance of our single-cell expression results, we performed pathway enrichment analysis^24, 25^ of DEG sets derived from each cell cluster along the crypt-villus axis. We first analyzed VSG-induced DEGs (filtered as previously described) to understand the effects of surgery alone on the epithelium. We then repeated these enrichment analyses with only genes “rescued” by VSG. In villus enterocytes, villus EECs, and some crypt-based goblet cells, genes downregulated by VSG were significantly enriched in pathways related to digestion and absorption of major macronutrients, vitamins, and minerals as well as cholesterol metabolism and PPAR signaling (**Figure 5A-B**). Of the VSG downregulated genes in the “rescue” category, digestion and absorption of fat and vitamins, cholesterol metabolism, and PPAR signaling remained top hits in most of the same villus clusters (**Figure 5A**). These findings suggest that VSG not only rescues HFD-induced defects in fat absorption and metabolism but also suppresses other macronutrient absorption pathways.

**Figure 5:**
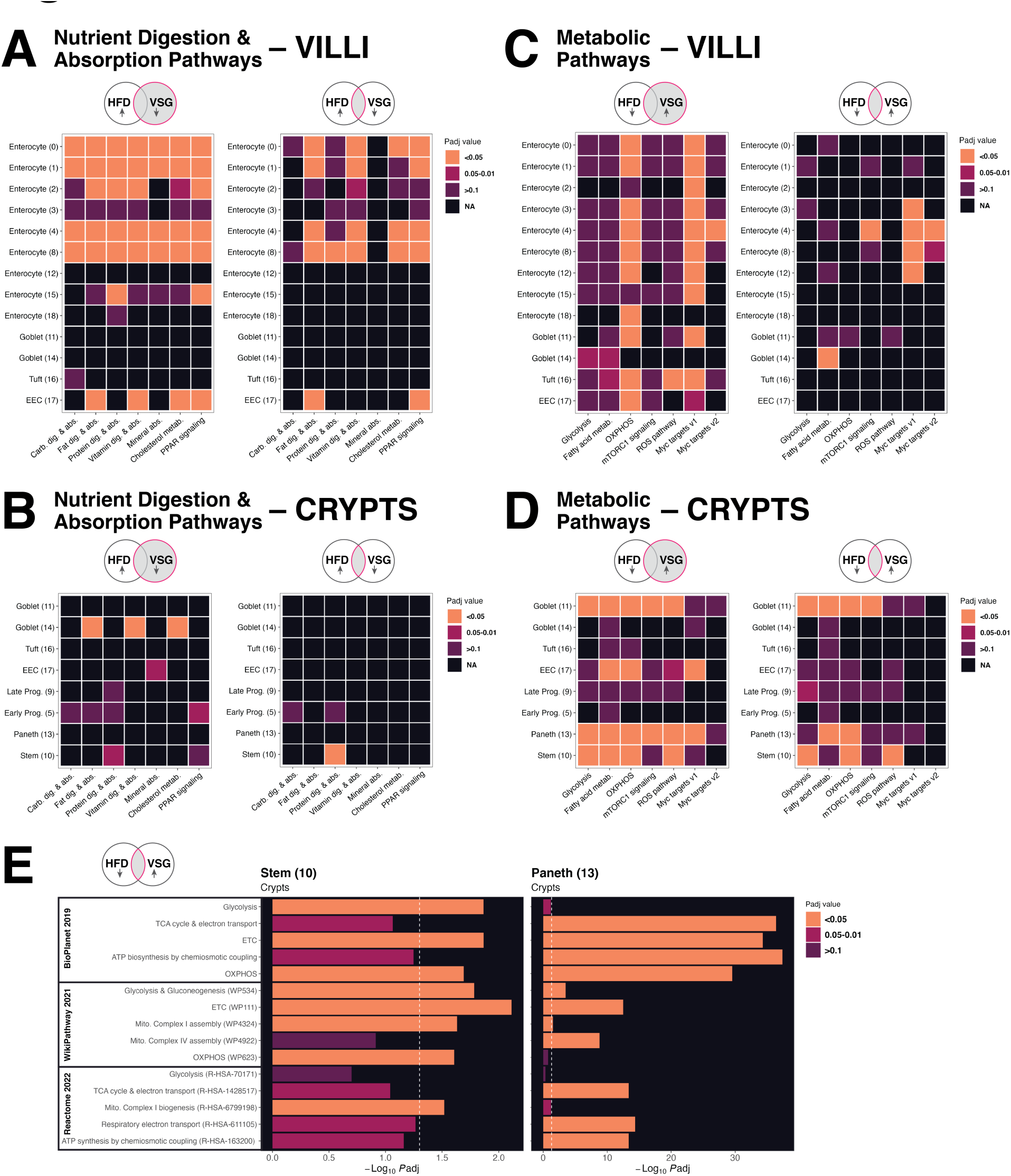
Nutrient absorption and metabolic function emerge as top pathway hits enriched among genes rescued by surgery. Heatmaps of pathway enrichment analysis results. Within each panel, the left plot shows results from genes differentially expressed by VSG, and the right plot shows the same analysis using genes rescued from HFD perturbation. Significant enrichment is marked by warmer colors among pathways relevant for nutrient digestion and absorption across (A) villus and (B) crypt clusters as well as metabolic pathways in (C) villus and (D) crypt clusters. (E) Bar plots of additional enrichment analysis results focused on metabolic pathways in the stem and Paneth clusters. *Padj* values are reported within the figure.

Conversely, genes upregulated by VSG were significantly enriched in a wide range of metabolic pathways. These included oxidative phosphorylation in crypt and villus clusters in addition to other crypt-based pathways such as glycolysis, fatty acid metabolism, and mTORC1 signaling (**Figure 5C-D**). Pathways pertaining to reactive oxygen species (ROS) and Myc targets also emerged as significant hits across crypt and villus lineages, specifically in stem and secretory cells for the former and both absorptive and secretory clusters for the latter (**Figure 5C-D**). Taken together, these results point to mitochondrial activity and biogenesis^26^ as possible mechanisms of VSG-induced metabolic improvements in the gut. Of the VSG upregulated genes in the “rescue” category, recovery of metabolic pathways was most pronounced in crypt-based goblet, stem, and Paneth cells (**Figure 5D**). Given the importance of metabolic regulation within the crypt to overall epithelial homeostasis^27–30^, we performed more expansive pathway enrichment analyses focused on stem and Paneth cells. We found that pathways related to the TCA cycle and electron transport chain, ATP biosynthesis, and mitochondrial complex assembly were among the most over-represented among genes rescued by VSG (**Figure 5E**). Notably, these results were based on nuclear-encoded genes, which prompted us to next explore changes in mitochondrial-encoded genes.

### VSG ameliorates defects in crypt-based mitochondrial gene expression induced by chronic HFD

To further investigate how HFD and VSG affect mitochondria within intestinal epithelial crypts, we compared the expression levels of all 13 mitochondrial protein-coding genes among our three experimental groups in both the stem and Paneth clusters. We observed perturbed expression of electron transport chain components with HFD as well as rescue of expression by VSG, especially among Complex I and Complex IV genes (**Figure 6A**). Several were rescued in both stem and Paneth clusters (e.g., *mt-Co1*), whereas others were more prominently rescued in one of the cell types (e.g., *mt-Nd6* in stem and *mt-Atp8* in Paneth).

**Figure 6:**
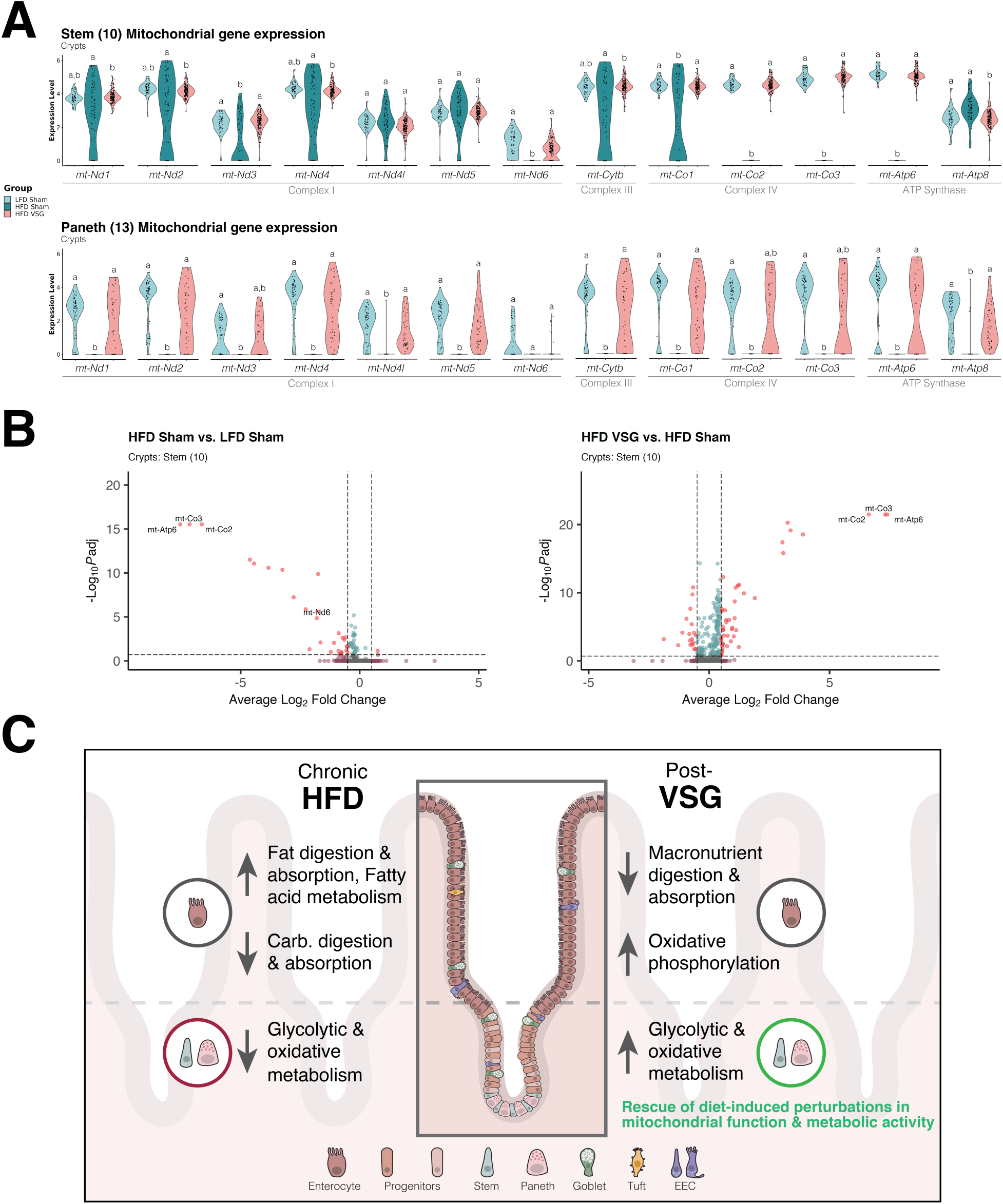
Vertical sleeve gastrectomy ameliorates changes in crypt-based mitochondrial gene expression driven by chronic high-fat diet. (A) Violin plots of stem and Paneth cell mitochondrial gene expression across experimental groups. Different letters indicate a statistically significant difference (*P* < 0.05 & *Padj* < 0.20) by MAST. (B) Volcano plots of significantly altered stem cell genes across dietary (left) and surgical (right) conditions, filtered by log2 fold change ±0.5 (vertical hashed lines) and *Padj* < 0.2 (horizontal hashed line). Notably, mitochondrially-encoded genes appear among the most differentially expressed even while accounting for the proportion of genes mapping to the mitochondrial genome as a covariate. (C) Proposed working model as to how chronic consumption of an obesogenic diet and treatment by VSG initiate gut adaptations through cell-type specific changes within the small intestinal epithelium.

Differences in cell and data quality can account for variance in the number of reads mapping to mitochondrial genes and introduce bias to our expression analyses. A stringent filtering threshold for mitochondrial reads (<20%) was applied across all single-cell datasets, which suggests that the aforementioned results are not likely caused by technical issues. Nonetheless, to make sure of this, we repeated the differential expression analysis while explicitly accounting for the per-cell proportion of reads mapping to the mitochondrial genome as a covariate. Indeed, we still found that mitochondrial protein-coding genes are among the most downregulated DEGs after HFD and the most upregulated after VSG (**Figure 6B**).

Altogether, our single-cell findings have led to the development of the following working model (**Figure 6C**), which proposes how the small intestinal epithelium adapts to HFD-induced obesity and treatment by VSG: (1) HFD elicits expression changes in villus enterocytes to accommodate increased fat digestion and metabolism at the expense of other macronutrients such as carbohydrates; (2) VSG reduces global macronutrient digestion and absorption in villus enterocytes and also programs them for increased oxidative phosphorylation; (3) among crypt-based stem and Paneth cells, nuclear-encoded genes involved in glycolysis, mitochondrial function, and metabolic activity are impaired by HFD and rescued by VSG; and (4) stem and Paneth mitochondrial-encoded genes (especially in the complex I and IV pathways) are among the most prominently rescued by VSG.

## Discussion

A variety of gut epithelial cell types sense and respond to environmental cues. Exposure to external factors, particularly dietary components, can adjust the balance of absorptive, defensive, and secretory functions distributed among these cells – either productively to maintain homeostasis or maladaptively to incite disease. In mice fed a diet high in fat and/or sugar, the intestinal epithelium shows a hyperproliferative response with lineage allocation skewed towards absorptive enterocytes at the expense of secretory lineages like EECs. These adaptations occur with both short- and long-term dietary interventions and correlate with defects in whole-body metabolism^5, 6^. From a therapeutic perspective, growing evidence suggests that the metabolic improvements observed after bariatric surgery arise from various changes in not only gut anatomy but also epithelial physiology^11^. Indeed, while post-surgical adaptations in gut endocrine signaling and their therapeutic effects continue to be explored, pharmacological approaches that recapitulate such outcomes have received increasing attention as attractive alternatives to treat metabolic disease^9^. Despite these advances, our understanding of how dietary and surgical interventions act through the gut to influence metabolic health remains limited.

To address this knowledge gap, we performed a high-resolution transcriptomic analysis of the small intestinal epithelium following VSG, one of the most common bariatric surgical procedures performed worldwide^12^. The current study builds on our previous efforts to characterize how VSG impacts epithelial differentiation via bulk RNA-sequencing of sorted intestinal stem cells^13^. Here, we leveraged single-cell technology to comprehensively survey the transcriptome of murine epithelial cells spanning the crypt-villus axis, both upon development of diet-induced obesity and after treatment by VSG. We identified all major epithelial lineages and revealed cell type-specific changes in gene expression between high- and low-fat diet-fed as well as VSG versus sham-operated mice. By comparing differential expression patterns resulting from these dietary and surgical interventions, we specified genes and pathways that VSG rescues from HFD perturbation and defined additional diet-independent changes of VSG. This drew our attention to crypt-based cell lineages, which showed greater proportions of rescue compared to villus cell types, particularly with genes downregulated by HFD and upregulated after VSG.

Given that epithelial differentiation and homeostasis are orchestrated by the crypt niche^20^, we further examined our single-cell results to determine how VSG-induced rescue of stem and Paneth cell genes might initiate adaptive changes within the gut. Several notable findings emerged. First, we distinguished patterns of rescued gene expression across different cell types (e.g., genes rescued in both stem and Paneth cells) in addition to expression rescued predominantly within specific lineages (e.g., genes rescued to a greater extent in either stem or Paneth cells). Among the latter group of genes, we expanded on a recent observation by Shin *et al*., who found that VSG increases expression of Reg3g, an antimicrobial peptide necessary to improve gut and metabolic function following dietary fiber supplementation and bariatric surgery^14^. While upregulation of *Reg3g* by VSG was observed broadly throughout the small intestine, we localized this effect mainly within Paneth cells and saw a concomitant trend toward rescued Paneth cell number via Lyz1 immunofluorescence. Next, we also noted crypt-centric rescue of genes relevant for a variety of metabolic pathways, such as glycolysis, oxidative phosphorylation, fatty acid metabolism, and mTORC1 signaling. Focusing on stem and Paneth cells, we noticed that VSG rescues both nuclear and mitochondrially encoded genes related to the electron transport chain, ATP biosynthesis, and assembly of mitochondrial complexes (especially complex I and IV). Altogether, our results suggest that chronic HFD impairs specific cell types along with the collective metabolic profile of the crypt niche and that these defects are ameliorated by VSG.

Previous studies have pointed to mitochondrial dysfunction within various tissues and organs, including the intestine^31^, the liver^32^, skeletal muscle^33^, and adipose depots^34, 35^, as potential contributors to metabolic disease pathogenesis. Bariatric surgery has also been shown to improve mitochondrial function in both preclinical models and patients^36–38^, though outside the context of the intestine. Our study emphasizes the underexplored role of intestinal epithelial cell metabolism in shaping the gut and overall metabolic health following dietary and surgical intervention. This coincides with a mounting body of evidence underscoring crypt metabolic activity as a crucial, conserved regulator of intestinal differentiation and homeostasis^29, 39, 40^. The discoveries in this area have highlighted the importance of stem cell mitochondrial respiration supported by glycolytic products (e.g., lactate) from Paneth cells^28, 30^. In line with this, we found that HFD reduces while VSG rescues Paneth cell expression of *Ldha*, which encodes a catalytic subunit of the glycolytic enzyme lactate dehydrogenase^23^. Furthermore, within stem cells we observed more prominent rescue of genes involved in mitigation of oxidative stress (e.g., *Gpx4*^21^) and apoptotic insults (e.g., *Prap1*^22^), which broadly pertain to mitochondrial function. These findings suggest that diet and bariatric surgery may induce gut adaptations through transcriptional changes that uniquely affect the metabolic profiles of different crypt-based lineages.

However, our data also prompted us to rethink the idea of strict stem- and Paneth-specific metabolic compartmentalization, as we noted more generalized rescue of other protein-coding components of oxidative metabolism and glycolysis across both cell types (e.g., *Cox7a2*, mitochondrially encoded electron transport chain genes, and *Gapdh*). Moreover, although mitochondrial respiration has been suggested as the main metabolic identity of stem cells^30^, a recent study has demonstrated the importance of glycolysis within *Lgr5*+ cells. Specifically, the authors showed that ablation of the glycolytic enzyme hexokinase 2 within this cell population was sufficient to perturb intestinal stem self-renewal and differentiation^41^. Another study used live-cell imaging to reveal a metabolic gradient that shifts from glycolysis to oxidative phosphorylation with cell proliferation and differentiation, respectively, along the crypt-villus axis^42^. Thus, a more nuanced perspective should be considered when weighing the relative contributions of different metabolic pathways and how they may dynamically impact the function of specific cell types over time. Future studies are needed to define how glycolytic and mitochondrial activity interact within and between different cell types to regulate overall intestinal homeostasis.

Beyond the cellular and transcriptomic changes observed in the crypts, diet- and surgery-induced adaptations in villi were mostly centered around enterocytic nutrient processing. VSG downregulated genes associated with macronutrient digestion and absorption, including those related to lipid handling and fat metabolism that were shown by us and others to be upregulated by HFD^5, 6^. At the same time, VSG increased expression of genes involved in oxidative phosphorylation across nearly all villus cell clusters, supporting the previously suggested metabolic trajectory of oxidative metabolism in mature, differentiated cells as opposed to glycolytic activity in proliferating, developing cells. These molecular changes coincided with subtle increases in crypt depth, villus height, and goblet cell number after VSG, which may reflect ways in which dietary and surgical interventions affect rates of epithelial differentiation and turnover through metabolic adjustments across cell lineages. Nonetheless, we acknowledge the need to interpret our findings carefully among several experimental factors, such as the specific mouse model used, the intestinal region interrogated, as well as the dietary composition and duration implemented during the study intervention. Other studies have drawn different conclusions. For example, while we saw no significant diet-induced changes in jejunal epithelial histomorphometry or secretory cell quantification here, others have reported increases, decreases, or no changes in these parameters, likely arising from differing study designs^5, 6, 43–45^. Therefore, our results do not definitively specify the effects of HFD and VSG on the gut but rather add to a growing picture of how the intestinal epithelium differentially responds to various contexts.

We recognize several limitations of our study. First, while there are known sex-dependencies in the development of diet-induced metabolic disease^46, 47^ as well as in treatment outcomes by bariatric surgery^16^, we focused only on male mice to minimize the impact of additional covariates in our single-cell analyses. Future studies should build on existing work^48^ to explore potential sex differences in gut adaptations initiated by diet and surgery. In addition, our single-cell analysis of certain cell types, such as EECs, was limited by their rarity in the epithelium. Follow up single-cell studies can leverage sorting or enrichment strategies to study changes more carefully in enteroendocrine gene expression and subtype profiling following bariatric surgery. Finally, we understand and appreciate that our findings conveyed here are restricted to the transcriptomic level of gene expression, which we hope will inspire future investigations into other aspects of gene regulation and protein function that are altered by VSG.

In conclusion, we present a comprehensive single-cell transcriptomic survey of the murine jejunal epithelium following treatment of diet-induced metabolic disease by VSG. We identified changes in gene expression within specific lineages throughout the crypt-villus axis, notably VSG-induced rescue of genes perturbed by HFD. Overall, our study contributes to resolving the potential cellular and molecular mechanisms that underlie gut adaptations and advances efforts to find more effective, less invasive ways to treat metabolic disease.

## Acknowledgements

The authors give their wholehearted thanks to the staff at the following core facilities for their invaluable assistance in this study: the Genomics and Microarray Core (Cancer Center Support Grant (P30CA046934), the Histology Core, and the Colorado Nutrition Obesity Research Center (NORC, DK048520) at the University of Colorado Anschutz Medical Campus; as well as the Animal Health Diagnostic Center Histology Core, the Institute of Biotechnology, and the Bioinformatics Facility at Cornell University. We also greatly appreciate our colleague, Dr. Matt Kanke, for his support and expertise in single-cell transcriptomic analyses as well as Danielle Leander for her surgical support. Figures were created with <u>BioRender.com</u> and Adobe Illustrator.

## Author Contributions

Conceptualization, K.K-L., K-S.K., D.A.S., and P.S.; Investigation, K.K-L., K-S.K., M.B., and K.F.; Formal Analysis, K.K-L. and K-S.K.; Visualization, K.K-L. and K-S.K.; Writing – Original Draft, K.K-L.; Writing – Review & Editing, K.K-L., K-S.K., M.B., K.F., D.A.S., and P.S.; Supervision, D.A.S., P.S.; Funding Acquisition, K.K-L., K-S.K., D.A.S., and P.S.

## Grants

This work was supported by the National Institutes of Health, National Institute of Diabetes and Digestive and Kidney Diseases (R01DK121995 and R01DK107282 to D. Sandoval, K01DK129367 to K.S. Kim), the NIH Office of Research Infrastructure Programs (F30OD031914 to K. Koch-Laskowski), as well as the American Diabetes Association (1-19-IBS-252 to D. Sandoval and 1-16-ACE-47 to P. Sethupathy).

## Declaration of Interests

The authors declare no competing interests.

## Methods

### In vivo studies

Five-week-old male C57BL/6J mice (n=20) were purchased from the Jackson Laboratory (Bar Harbor, ME, USA) and were individually housed in a 12-hour light/dark cycle environment with ad libitum access to water and food. The animal room was maintained at a temperature of 25°C with 50%-60% humidity. Following an acclimation period, mice were assigned to receive either a 10% LFD (Research Diet; catalog D12450J) or a 60% HFD (Research Diet; catalog D12492) for a duration of 16 weeks before surgical intervention.

Mice were matched for body weight and body fat, then subjected to sham or VSG surgery as previously described^15^. Briefly, mice fed a 60% HFD (n=5) were anesthetized, and a small laparotomy incision was made in the abdominal wall. The lateral 80% of the stomach along the greater curvature was excised, and sleeve was created using a simple continuous suture (8-0 Prolene). Simple interrupted sutures were occasionally utilized to reinforce the strength of the sleeve. Sham surgery was performed on LFD-fed mice (n=6) and HFD-fed mice (n=6) by applying gentle pressure on the stomach with blunt forceps. During the initial 3 days following surgery, the animals were fed a DietGel^®^ boost (Clear H2O Inc, Westbrook, ME, USA) and subsequently returned to their original LFD or HFD.

Body weight and food intake were monitored for 5 weeks post-surgery. Body composition was assessed before and 5 weeks after surgery using an EchoMRI instrument (EchoMRI LLC, Houston, TX, USA). At the 4-week mark post-surgery, an oral glucose tolerance test (OGTT) was conducted after a 5- to 6-hour fast, orally administering a 2 g/kg dose of a 50% dextrose solution.

All animal studies were performed according to an approved protocol by the Institutional Animal Care and Use Committee (IACUC) at the University of Colorado Anschutz Medical Campus as well as protocols outlined in the National Institutes of Health (NIH) guide for the care and use of laboratory animals (NIH Publications No. 8023, revised 1978).

### Small intestinal crypt/villus collections and single-cell isolations

Five weeks post-surgery, overnight-fasted mice were euthanized via CO_2_ inhalation. The abdominal cavity was promptly opened, and the small intestine was collected and divided into three segments (duodenum, jejunum and ileum). Subsequently, the segments were flushed with ice-cold PBS (Gibco, ThermoFisher, Waltham, MA, USA) to remove luminal contents. A small portion of each segment (∼0.5 cm) was isolated for histologic analysis. Each jejunal segment was opened longitudinally and placed in individual tubes containing cold DMEM (ThermoFisher) before immediate processing with a solution of 3mM EDTA (Sigma-Aldrich, St. Louis, MO, USA) in PBS for cell dissociation. During processing in the EDTA/PBS solution, the intestinal segments were manually scrapped and subsequently filtered through a 70 µm cell strainer to separate crypts from villi.

For single-cell dissociation, the crypts and villi were resuspended separately in a cold solution of 0.04% bovine serum albumin (BSA)/PBS, followed by processing with 0.3 U/ml dispase/HBSS, DNAse1/FBS, and 0.04% BSA/PBS solutions (all reagents purchased from Sigma-Aldrich). The viability of and number of dissociated crypt and villus cells were measured using the ThermoFisher ViCell counter (ThermoFisher), with an average viability of 71% for crypts and 66% for villus cells. Approximately 3.0 × 10^7^ crypt cells and 3.5 × 10^7^ villus cells from each sample were pooled (n=3 for LFD-sham and HFD-sham, n=2 for HFD-VSG) and diluted to a concentration of 1000 cells/μL.

### Single-cell library preparation and sequencing

Single-cell RNA-sequencing library preparation was performed by the Genomics and Microarray Core at the University of Colorado Anschutz Medical Campus. Using the 10x Genomics Chromium Next GEM 3’ v3.1 kit, single-cell suspensions were processed by loading roughly ∼16,500 cells to capture a target number of 10,000 per sample with which to generate libraries. Sequencing was run on the NovaSeq 6000 platform (Illumina, San Diego, CA, USA) to obtain at least 50,000 reads per cell.

### Single-cell transcriptomic analysis pipeline

FASTQ files were generated from the raw single-cell sequencing data and aligned to the mouse genome (mm10) via CellRanger (v.6.0.0). To help ensure that our transcriptomic signal originated from productive gel beads-in-emulsion rather than ambient cell-free transcripts, raw and filtered gene/count matrices produced by CellRanger were processed through SoupX^49^ via the default parameters of the autoEstCont method. The estimated background contamination fractions (rho) for each sample were as follows: LFD sham crypts = 0.030, LFD sham villi = 0.046, HFD sham crypts = 0.279, HFD sham villi = 0.165, HFD VSG crypts = 0.035, HFD VSG villi = 0.021. Corrected matrices with ambient RNA removed were initialized in Seurat (v4.1)^50^ for all downstream quality control and analysis steps. First, multiplets were computationally accounted for and removed via scDlbFinder^51^. Next, low-quality cells with less than 750 genes detected or greater than 20% of reads mapping to mitochondrial genes were filtered out. With the remaining high-quality cells, samples were normalized and integrated using the standard Seurat SCTransform (v2)^52, 53^ workflow based on the 2000 most variable genes while controlling for the number of genes per cell and the proportion of mitochondrial reads mapped. Subsequent clustering visualizations projected via UMAP were derived from 30 principal component analysis dimensions at a resolution of 0.4 and later customized to different color palettes using ggplot2. Highly enriched genes in each cluster were determined by the FindAllMarkers function in Seurat, which identified the most upregulated genes compared to all other clusters; marker enrichment was defined as having a log fold change of at least 0.50 via MAST^54^ with per-cell gene number and mitochondrial reads as covariates. Cell types were assigned to each cluster based on overlap of enriched genes with known markers of each intestinal epithelial lineage as annotated previously by Haber et al^3^. Additional overlap with immune cell markers (e.g., *Gzma*, *Gzmb*, *Itgae*, *Ccl5*, *Cd7*, *Cd69*, *Cd3g*, *Cd8a*) identified two immune clusters that were removed from subsequent analyses. Focusing on epithelial clusters, cells were assigned a cell-cycle score based on expression of G2/M and S phase markers via Seurat’s CellCycleScoring function. Pseudotime analysis was performed on Seurat-defined clusters via Monocle3^55^ with root nodes originating in the stem cell cluster.

Within each intestinal epithelial cell cluster across the crypt-villus axis, differential expression analysis between experimental groups (i.e., HFD VSG vs. HFD Sham, HFD Sham vs. LFD Sham, and HFD VSG vs. LFD Sham) was performed using the FindMarkers function in Seurat; significant differential expression of genes was determined via MAST^54^. Changes in mitochondrial gene expression were confirmed in a repeated analysis accounting for genes per cell and percentage of mitochondrial reads as covariates. Differentially expressed genes (DEGs) from each cluster were filtered (*P* < 0.05, *P*adj < 0.20, log2FC > |0.50|) and subsequently compared by diet (i.e., HFD sham vs. LFD sham) and surgery (i.e., HFD VSG vs. HFD sham). These DEGs were further analyzed in a cluster-specific manner via a Venn diagram approach to identify genes demonstrating reciprocal changes in expression overlapped between dietary and surgical comparisons (see **Figure 4A**).

Pathway enrichment analysis was performed by inputting filtered DEG sets (*P* < 0.05, *P*adj < 0.20, log2FC > |0.50|) into Enrichr^24, 25^, a comprehensive online database of gene set annotations and libraries. Adjusted *P* values for enrichment terms were manually collected and projected by heatmap using ggplot2. Nutrient digestion and absorption pathways were derived strictly from Enrichr’s KEGG Human 2021 database. Metabolic pathways were initially surveyed across all clusters via Enrichr’s Molecular Signatures Database Hallmark 2020; further investigation of metabolic pathways in the stem and Paneth clusters leveraged the BioPlanet 2019, WikiPathway 2021, and Reactome 2022 databases.

### Histologic analysis

A small portion from each intestinal segment was fixed in 10% neutral-buffered formalin for 24 hours and submitted to the University of Colorado Histology Shared Resource for paraffin embedding and slide preparation. All images were captured using a BX53 Olympus scope. Morphometric analyses were performed by analyzing hematoxylin and eosin-stained sections in ImageJ. Measurement of crypt depth and villus height spanned from the crypt base to the top of the transit-amplifying zone then to the villus tip, respectively. Averaged morphometric values were based upon inclusion of at least ten intact crypts and villi per animal in each experimental group. Using Alcian blue-stained sections, goblet cells were quantified in both crypts and villi through manual counting of positively labeled cells situated along the outline of the intestinal epithelium. Alcian blue counts were averaged per animal within crypts and villi separately. Paneth cell quantification was assessed via immunofluorescent staining for Lyz1. Here, sections were deparaffinized using a series of xylene and ethanol solutions, rehydrated in cold water, and permeabilized with methanol. After boiling in antigen retrieval solution (10mM citric acid, 0.05% v/v Tween-20, pH = 6.0) for 20 minutes, sections were blocked with 10% v/v normal goat serum in PBS for 1 hour. Sections were incubated overnight with primary antibody (rabbit anti-Lyz1, diluted 1:1000 in PBS with 1% w/v bovine serum albumin) at 4°C followed by a 1-hour room temperature incubation with secondary antibody (goat anti-rabbit Alexa Fluor 594 diluted 1:1000 in PBS with 1%w/v bovine serum albumin). Nuclei were subsequently counterstained with DAPI (diluted 1:1000 in PBS). Lyz1+ cells were manually quantified in intact crypts and averaged per animal.

### Statistics

Statistical analyses of *in vivo* data were performed using GraphPad Prism 9 (GraphPad Software, San Diego, CA, USA). Body weight/composition, oral glucose tolerance testing, and histological data were analyzed using ordinary one- or two-way ANOVA, as applicable, to determine significant main effects and interactions between independent variables. Food intake measurements were assessed through mixed-effects analysis. Significant differences (*P*adj < 0.05) were determined by Tukey’s post hoc testing. Data are presented as mean ± SEM. Single-cell RNA-sequencing analyses were performed using R. Differential gene expression outcomes were determined via MAST^54^ and were considered significant with *P* < 0.05 and *P*adj < 0.20.

## Notes

### Competing Interest Statement

The authors have declared no competing interest.

## References

1. Beumer, J. & Clevers, H. Cell fate specification and differentiation in the adult mammalian intestine. Nat. Rev. Mol. Cell Biol. 22, 39–53 (2021).

2. Gribble, F. M. & Reimann, F. Function and mechanisms of enteroendocrine cells and gut hormones in metabolism. Nat. Rev. Endocrinol. 15, 226–237 (2019).

3. Haber, A. L. et al. A single-cell survey of the small intestinal epithelium. Nature 551, 333– 339 (2017).

4. Gehart, H. et al. Identification of Enteroendocrine Regulators by Real-Time Single-Cell Differentiation Mapping. Cell 176, 1158–1173.e16 (2019).

5. Enriquez, J. R. et al. A dietary change to a high-fat diet initiates a rapid adaptation of the intestine. Cell Rep. 41, 111641 (2022).

6. Aliluev, A. Diet-induced alteration of intestinal stem cell function underlies obesity and prediabetes in mice. 3, 35 (2021).

7. Heymsfield, S. B. & Wadden, T. A. Mechanisms, Pathophysiology, and Management of Obesity. N. Engl. J. Med. 376, 254–266 (2017).

8. Blüher, M. Obesity: global epidemiology and pathogenesis. Nat. Rev. Endocrinol. 15, 288– 298 (2019).

9. Gimeno, R. E., Briere, D. A. & Seeley, R. J. Leveraging the Gut to Treat Metabolic Disease. Cell Metab. 31, 679–698 (2020).

10. Kim, K.-S. & Sandoval, D. A. Endocrine Function after Bariatric Surgery. in Comprehensive Physiology 783–798 (American Cancer Society, 2017). doi:10.1002/cphy.c160019.

11. Seeley, R. J., Chambers, A. P. & Sandoval, D. A. The role of gut adaptation in the potent effects of multiple bariatric surgeries on obesity and diabetes. Cell Metab. 21, 369–378 (2015).

12. Welbourn, R. et al. Bariatric Surgery Worldwide: Baseline Demographic Description and One-Year Outcomes from the Fourth IFSO Global Registry Report 2018. Obes. Surg. 29, 782–795 (2019).

13. Kim, K.-S. et al. Vertical sleeve gastrectomy induces enteroendocrine cell differentiation of intestinal stem cells through bile acid signaling. JCI Insight 7, e154302 (2022).

14. Shin, J. H. et al. The gut peptide Reg3g links the small intestine microbiome to the regulation of energy balance, glucose levels, and gut function. Cell Metab. 34, 1765–1778.e6 (2022).

15. Kim, K.-S. et al. Vertical sleeve gastrectomy induces enteroendocrine cell differentiation of intestinal stem cells through bile acid signaling. JCI Insight (2022) doi:10.1172/jci.insight.154302.

16. Hutch, C. R. et al. Diet-Dependent Sex Differences in the Response to Vertical Sleeve Gastrectomy. Am. J. Physiol.-Endocrinol. Metab. (2021) doi:10.1152/ajpendo.00060.2021.

17. Schauer, P. R. et al. Bariatric Surgery versus Intensive Medical Therapy in Obese Patients with Diabetes. http://dx.doi.org/10.1056/NEJMoa1200225 https://www.nejm.org/doi/10.1056/NEJMoa1200225 (2012) doi:10.1056/NEJMoa1200225.

18. Evers, S. S. et al. Continuous glucose monitoring reveals glycemic variability and hypoglycemia after vertical sleeve gastrectomy in rats. Mol. Metab. 32, 148–159 (2020).

19. Stefater, M. A. et al. Sleeve Gastrectomy Induces Loss of Weight and Fat Mass in Obese Rats, but Does Not Affect Leptin Sensitivity. Gastroenterology 138, 2426–2436.e3 (2010).

20. Gehart, H. & Clevers, H. Tales from the crypt: new insights into intestinal stem cells. Nat. Rev. Gastroenterol. Hepatol. 16, 19–34 (2019).

21. Mayr, L. et al. Dietary lipids fuel GPX4-restricted enteritis resembling Crohn’s disease. Nat. Commun. 11, 1775 (2020).

22. Wolfarth, A. A. et al. Proline-Rich Acidic Protein 1 (PRAP1) Protects the Gastrointestinal Epithelium From Irradiation-Induced Apoptosis. Cell. Mol. Gastroenterol. Hepatol. 10, 713– 727 (2020).

23. Wu, Q. et al. Intestinal hypoxia-inducible factor 2α regulates lactate levels to shape the gut microbiome and alter thermogenesis. Cell Metab. 33, 1988–2003.e7 (2021).

24. Chen, E. Y. et al. Enrichr: interactive and collaborative HTML5 gene list enrichment analysis tool. BMC Bioinformatics 14, 128 (2013).

25. Kuleshov, M. V. et al. Enrichr: a comprehensive gene set enrichment analysis web server 2016 update. Nucleic Acids Res. 44, W90–97 (2016).

26. Morrish, F. & Hockenbery, D. MYC and Mitochondrial Biogenesis. Cold Spring Harb. Perspect. Med. 4, a014225–a014225 (2014).

27. Urbauer, E., Rath, E. & Haller, D. Mitochondrial Metabolism in the Intestinal Stem Cell Niche—Sensing and Signaling in Health and Disease. Front. Cell Dev. Biol. 8, 602814 (2021).

28. Ludikhuize, M. C. et al. Mitochondria Define Intestinal Stem Cell Differentiation Downstream of a FOXO/Notch Axis. Cell Metab. 32, 889–900.e7 (2020).

29. Rath, E., Moschetta, A. & Haller, D. Mitochondrial function — gatekeeper of intestinal epithelial cell homeostasis. Nat. Rev. Gastroenterol. Hepatol. 15, 497–516 (2018).

30. Rodríguez-Colman, M. J. et al. Interplay between metabolic identities in the intestinal crypt supports stem cell function. Nature 543, 424–427 (2017).

31. Guerbette, T., Boudry, G. & Lan, A. Mitochondrial function in intestinal epithelium homeostasis and modulation in diet-induced obesity. Mol. Metab. 63, 101546 (2022).

32. Fromenty, B. & Roden, M. Mitochondrial alterations in fatty liver diseases. J. Hepatol. 78, 415–429 (2023).

33. Bakkman, L. et al. Reduced Respiratory Capacity in Muscle Mitochondria of Obese Subjects. Obes. Facts 3, 1–1 (2010).

34. Da Eira, D., Jani, S. & Ceddia, R. B. An obesogenic diet impairs uncoupled substrate oxidation and promotes whitening of the brown adipose tissue in rats. J. Physiol. 601, 69–82 (2023).

35. AlZaim, I., Eid, A. H., Abd-Elrahman, K. S. & El-Yazbi, A. F. Adipose tissue mitochondrial dysfunction and cardiometabolic diseases: On the search for novel molecular targets. Biochem. Pharmacol. 206, 115337 (2022).

36. Sacks, J. et al. Effect of Roux-en-Y gastric bypass on liver mitochondrial dynamics in a rat model of obesity. Physiol. Rep. 6, e13600 (2018).

37. Fernström, M. et al. Improved Muscle Mitochondrial Capacity Following Gastric Bypass Surgery in Obese Subjects. Obes. Surg. 26, 1391–1397 (2016).

38. Nijhawan, S., Richards, W., O’Hea, M. F., Audia, J. P. & Alvarez, D. F. Bariatric surgery rapidly improves mitochondrial respiration in morbidly obese patients. Surg. Endosc. 27, 4569–4573 (2013).

39. McCauley, H. A. et al. Enteroendocrine Cells Protect the Stem Cell Niche by Regulating Crypt Metabolism in Response to Nutrients. Cell. Mol. Gastroenterol. Hepatol. S2352345X22002673 (2023) doi:10.1016/j.jcmgh.2022.12.016.

40. Deng, H., Takashima, S., Paul, M., Guo, M. & Hartenstein, V. Mitochondrial dynamics regulates Drosophila intestinal stem cell differentiation. Cell Death Discov. 4, 81 (2018).

41. Li, C. et al. Glycolytic Regulation of Intestinal Stem Cell Self-Renewal and Differentiation. Cell. Mol. Gastroenterol. Hepatol. 15, 931–947 (2023).

42. Stringari, C. et al. Metabolic trajectory of cellular differentiation in small intestine by Phasor Fluorescence Lifetime Microscopy of NADH. Sci. Rep. 2, 568 (2012).

43. Soares, A., Beraldi, E. J., Ferreira, P. E. B., Bazotte, R. B. & Buttow, N. C. Intestinal and neuronal myenteric adaptations in the small intestine induced by a high-fat diet in mice. BMC Gastroenterol. 15, 3 (2015).

44. Mah, A. T., Van Landeghem, L., Gavin, H. E., Magness, S. T. & Lund, P. K. Impact of diet-induced obesity on intestinal stem cells: hyperproliferation but impaired intrinsic function that requires insulin/IGF1. Endocrinology 155, 3302–3314 (2014).

45. Beyaz, S. et al. High-fat diet enhances stemness and tumorigenicity of intestinal progenitors. Nature 531, 53–58 (2016).

46. Huang, K.-P. et al. Sex differences in response to short-term high fat diet in mice. Physiol. Behav. 221, 112894 (2020).

47. Millward, D. J. et al. Sex differences in the composition of weight gain and loss in overweight and obese adults. Br. J. Nutr. 111, 933–943 (2014).

48. Zhou, W., Davis, E. A., Li, K., Nowak, R. A. & Dailey, M. J. Sex differences influence intestinal epithelial stem cell proliferation independent of obesity. Physiol. Rep. 6, e13746 (2018).

49. Young, M. D. & Behjati, S. SoupX removes ambient RNA contamination from droplet-based single-cell RNA sequencing data. GigaScience 9, giaa151 (2020).

50. Hao, Y. et al. Integrated analysis of multimodal single-cell data. Cell 184, 3573–3587.e29 (2021).

51. Germain, P.-L., Lun, A., Meixide, C. G., Macnair, W. & Robinson, M. D. Doublet identification in single-cell sequencing data. (2022).

52. Hafemeister, C. & Satija, R. Normalization and variance stabilization of single-cell RNA-seq data using regularized negative binomial regression. Genome Biol. 20, 296 (2019).

53. Choudhary, S. & Satija, R. Comparison and evaluation of statistical error models for scRNA-seq. Genome Biol. 23, 27 (2022).

54. Finak, G. et al. MAST: a flexible statistical framework for assessing transcriptional changes and characterizing heterogeneity in single-cell RNA sequencing data. Genome Biol. 16, 278 (2015).

55. Trapnell, C. et al. The dynamics and regulators of cell fate decisions are revealed by pseudotemporal ordering of single cells. Nat. Biotechnol. 32, 381–386 (2014).

