## Supplemental Figures for "Intestinal epithelial adaptations to vertical sleeve gastrectomy defined at single-cell resolution"

### Supplemental Figure 1

## A

**CRYPTS** **VILLI**

**LFD Sham**

**HFD Sham**

**HFD VSG**

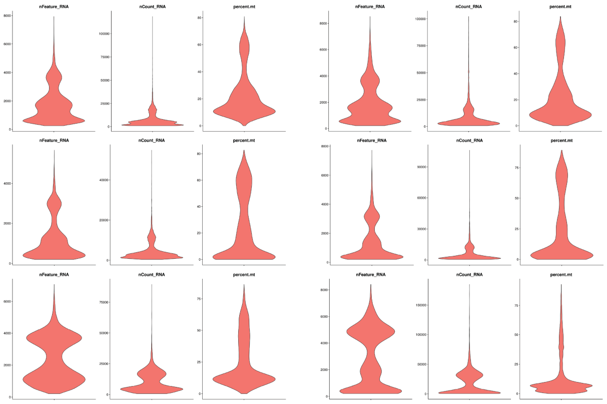


## B

**Quality Control Filtering Thresholds:** nFeature > 750 & percent.mt < 20%

| Sample | # Cells: Pre-Filtering | # Cells: Post-Filtering | # Cells: Immune removed |
|---|---|---|---|
| LFD Sham Crypts | 9654 | 4168 | 3637 |
| LFD Sham Villi | 8733 | 4251 | 3423 |
| HFD Sham Crypts | 8374 | 3369 | 3232 |
| HFD Sham Villi | 10162 | 3938 | 3736 |
| HFD VSG Crypts | 8999 | 4708 | 4326 |
| HFD VSG Villi | 6972 | 4077 | 3490 |
| Total | 52,894 | 24,511 | 21,844 |

| Metadata | # Cells: Pre-Filtering | # Cells: Post-Filtering | # Cells: Immune removed |
|---|---|---|---|
| LFD | 18,387 | 8419 | 7060 |
| HFD | 34,507 | 16,092 | 14,784 |

| Metadata | # Cells: Pre-Filtering | # Cells: Post-Filtering | # Cells: Immune removed |
|---|---|---|---|
| Sham | 36,923 | 15,726 | 14,028 |
| VSG | 15,971 | 8785 | 7816 |

| Metadata | # Cells: Pre-Filtering | # Cells: Post-Filtering | # Cells: Immune removed |
|---|---|---|---|
| Crypts | 27,027 | 12,245 | 11,195 |
| Villi | 25,867 | 12,266 | 10,649 |

## C

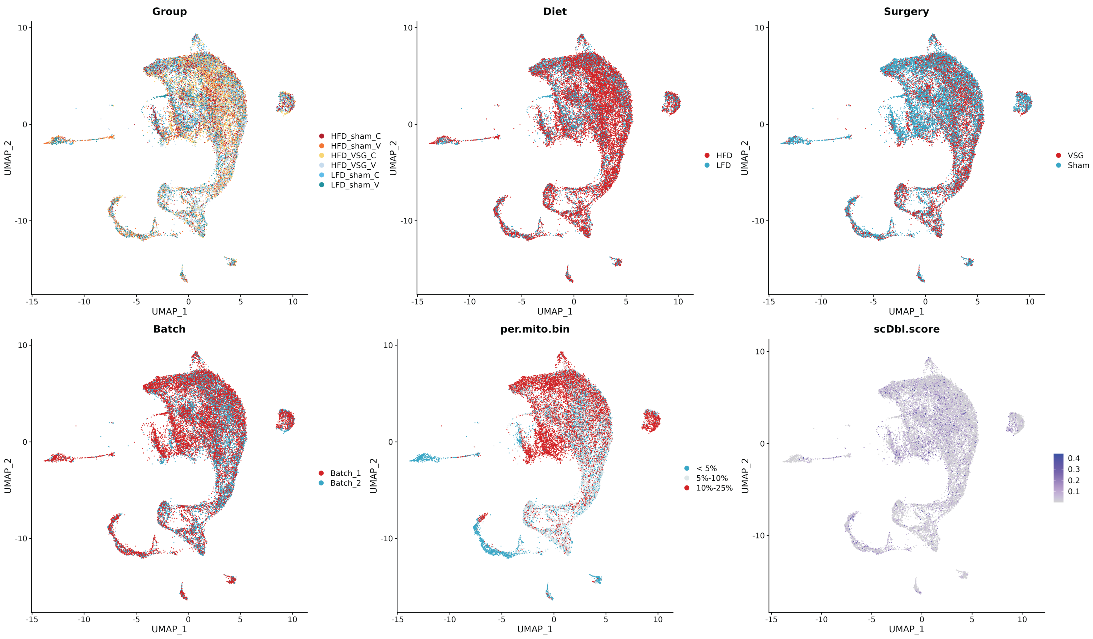

Working title: **Intestinal epithelial adaptations to vertical sleeve gastrectomy defined at single-cell resolution**

**Supplementary Figure 1: Single-cell transcriptomic quality control outcomes**

(A) Violin plots of the number of genes per cell (nFeature_RNA), the number of reads detected by unique molecular identifier (nCount_RNA), and the percentage of genes per cell that map to the mitochondrial genome (percent.mt). Each of these quality control metrics is shown for cell samples derived from the crypts or villi of each experimental group.

(B) Tables of the number of cells from each sample group and experimental condition before and after quality control filtering. Values in the rightmost column represent the finalized number of cells used in the present analyses.

(C) Uniform Manifold Approximation and Projection overlaps of various metadata, including sample group, dietary condition, surgical treatment, sample isolation batch, percentage bins of mitochondrially mapped genes, and doublet detection by scDblFinder.

Working title: **Intestinal epithelial adaptations to vertical sleeve gastrectomy defined at single-cell resolution**

### Supplemental Figure 2

#### VILLI

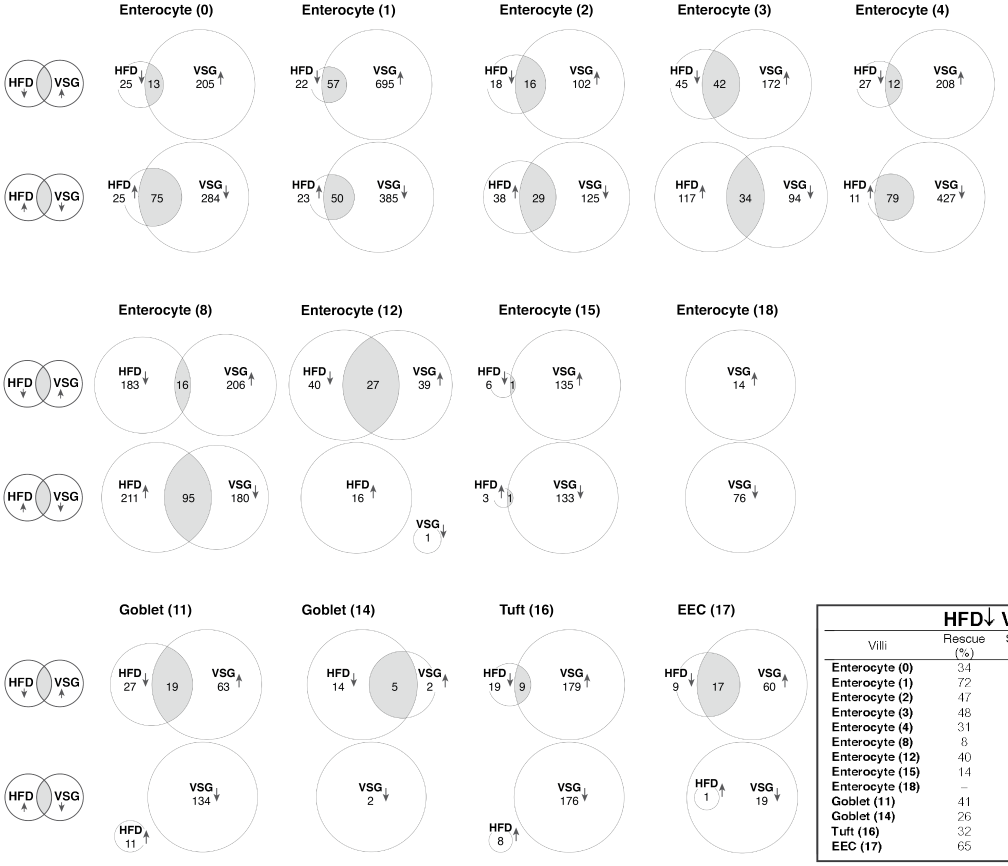


| | HFD↓ VSG↑ | | HFD↑ VSG↓ | |
|---|---|---|---|---|
| Villi | Rescue (%) | Specificity (%) | Rescue (%) | Specificity (%) |
| **Enterocyte (0)** | 34 | 6 | 75 | 21 |
| **Enterocyte (1)** | 72 | 8 | 68 | 11 |
| **Enterocyte (2)** | 47 | 14 | 43 | 19 |
| **Enterocyte (3)** | 48 | 20 | 23 | 27 |
| **Enterocyte (4)** | 31 | 5 | 88 | 16 |
| **Enterocyte (8)** | 8 | 7 | 31 | 35 |
| **Enterocyte (12)** | 40 | 41 | 0 | 0 |
| **Enterocyte (15)** | 14 | 0.7 | 25 | 0.7 |
| **Enterocyte (18)** | – | 0 | – | 0 |
| **Goblet (11)** | 41 | 23 | 0 | 0 |
| **Goblet (14)** | 26 | 71 | – | 0 |
| **Tuft (16)** | 32 | 5 | 0 | 0 |
| **EEC (17)** | 65 | 22 | 100 | 5 |

#### CRYPTS

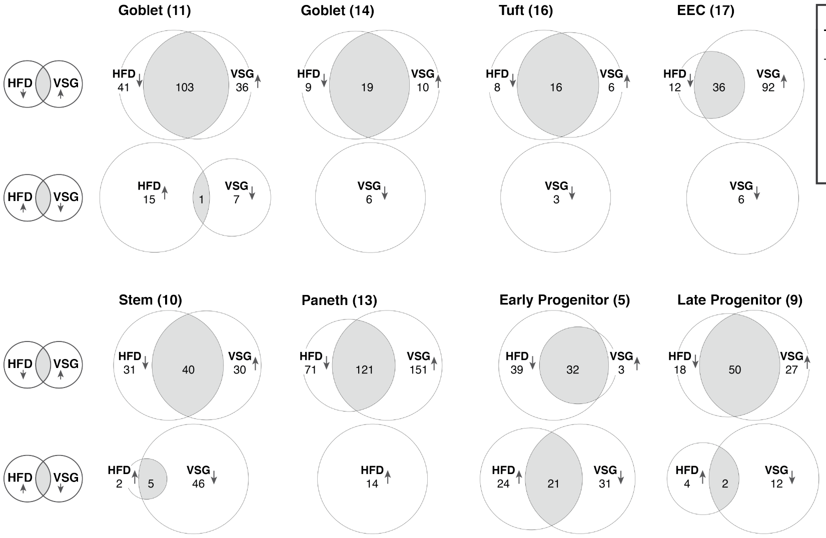


| | HFD↓ VSG↑ | | HFD↑ VSG↓ | |
|---|---|---|---|---|
| Crypts | Rescue (%) | Specificity (%) | Rescue (%) | Specificity (%) |
| **Goblet (11)** | 72 | 74 | 6 | 13 |
| **Goblet (14)** | 68 | 66 | – | 0 |
| **Tuft (16)** | 67 | 73 | – | 0 |
| **EEC (17)** | 75 | 28 | – | 0 |
| **Late Prog. (9)** | 74 | 65 | 33 | 14 |
| **Early Prog. (5)** | 45 | 91 | 47 | 40 |
| **Paneth (13)** | 63 | 44 | 0 | – |
| **Stem (10)** | 56 | 57 | 71 | 10 |

Working title: **Intestinal epithelial adaptations to vertical sleeve gastrectomy defined at single-cell resolution**

**Supplementary Figure 2: Comprehensive analysis of rescue and specificity across all intestinal epithelial cell clusters**

Venn diagrams of the number of differentially expressed genes identified between dietary and surgical comparisons across all crypt and villus clusters. Each cell cluster has two diagrams to illustrate both comparisons of differential expression (i.e., genes downregulated by HFD and upregulated by VSG, and vice versa). In some cases, there were no differentially expressed genes detected within a certain condition or shared between conditions of a given cell cluster, in which case the Venn diagram is altered appropriately. These values were used to calculate the proportions of rescue and specificity within each cell type, which are included in the accompanying tables.

Working title: **Intestinal epithelial adaptations to vertical sleeve gastrectomy defined at single-cell resolution**

### Supplemental Figure 3

**A**

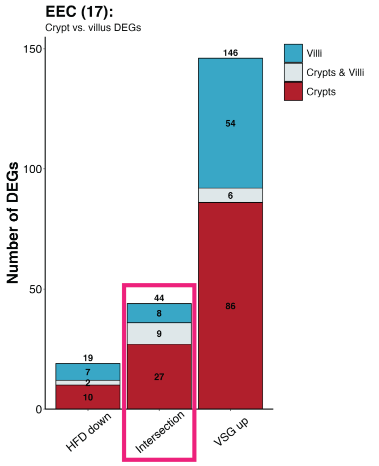


**B**

#### EEC Rescued Genes

| Crypts | | Crypts & Villi | Villi |
|---|---|---|---|
| Aldob | Guk1 | Defa24 | Defa17 |
| Apoa1 | Ndufb7 | mt-Atp6 | Defa21 |
| Apoa4 | Prap1 | mt-Co1 | Defa22 |
| Apoc2 | Prelid1 | mt-Co2 | Defa30 |
| Apoc3 | Rbp2 | mt-Co3 | Itln1 |
| Ckmt1 | Reg3b | mt-Cytb | Lyz1 |
| Crip1 | Rpl28 | mt-Nd2 | mt-Nd1 |
| Dbi | Rps12 | mt-Nd3 | Spink4 |
| Fabp1 | Serf2 | Zg16 | |
| Fabp2 | Spink1 | | |
| Fth1 | Ssr4 | | |
| Gpx4 | Vamp8 | | |
| Gsto1 | 2200002D01Rik | | |
| Guca2b | | | |

**C**

#### CRYPTS

**Stem (10)**

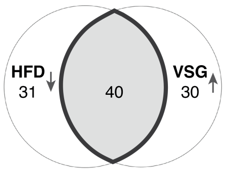


**Rescue:** 56%
**Specificity:** 57%

**Paneth (13)**

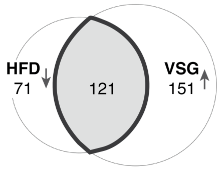


**Rescue:** 63%
**Specificity:** 44%

**Rescued genes: HFD↓ & VSG↑**

| Stem 21 | Stem & Paneth 19 | Paneth 102 | | |
|---|---|---|---|---|
| 2200002D01Rik | Actb | Actn4 | Cox8a | mt-Nd5 |
| Abhd2 | Aldob | Anxa4 | Ctnnb1 | Myl6 |
| Apoc3 | Apoa1 | Arpc2 | Cyb5r3 | Myo1a |
| Atp5md | Apoa4 | Atp1a1 | Cycs | Ndufa13 |
| Crip1 | Apoc2 | Atp1b1 | Dpp4 | Ndufb9 |
| Defa24 | AY036118 | Atp5a1 | Epcam | Oaz1 |
| Defa30 | B2m | Atp5b | Espn | Pfn1 |
| Fabp1 | Cox7a2 | Atp5c1 | Etfb | Pgrmc1 |
| Gpx4 | Dbi | Atp5d | Ezr | Phgr1 |
| Lct | Fabp2 | Atp5e | Fau | Pigr |
| Mgst3 | Fth1 | Atp5f1 | Ftl1 | Prdx1 |
| mt-Co3 | Gapdh | Atp5g1 | Gm42418 | Reg3g |
| mt-Nd6 | Guca2b | Atp5g3 | Gsto1 | Rfk |
| Ndufa2 | Ifi27l2b | Atp5h | H2-D1 | Rplp1 |
| Plac8 | mt-Atp6 | Atp5j | H2-K1 | Rps14 |
| Prap1 | mt-Co2 | Atp5j2 | H2-Q2 | Rps26 |
| Smim24 | Ndufa6 | Atp5l | Hadh | Scp2 |
| Spink1 | Selenop | Atp5o | Hadha | Serpinb6a |
| Tff3 | Serf2 | Calm1 | Hint1 | Slc25a3 |
| Tm4sf5 | | Calm3 | Krt8 | Slc25a5 |
| Zg16 | | Calml4 | Lap3 | Slc43a2 |
| | | Cdh17 | Ldha | Slc5a1 |
| | | Cdhr2 | Lgals4 | Smdt1 |
| | | Cdhr5 | Marc2 | Tle5 |
| | | Cfl1 | Mdh1 | Tma7 |
| | | Chchd2 | Mdh2 | Tmbim6 |
| | | Cldn7 | mt-Atp8 | Tpm1 |
| | | Clk1 | mt-Co1 | Txn1 |
| | | Cox4i1 | mt-Cytb | Txndc17 |
| | | Cox5a | mt-Nd1 | Ubb |
| | | Cox6b1 | mt-Nd2 | Uqcrb |
| | | Cox6c | mt-Nd3 | Uqcrh |
| | | Cox7b | mt-Nd4 | Uqcrq |
| | | Cox7c | mt-Nd4l | Vil1 |

Working title: **Intestinal epithelial adaptations to vertical sleeve gastrectomy defined at single-cell resolution**

**Supplementary Figure 3: Cell type-specific rescue of genes downregulated by HFD and upregulated by VSG**
(A) Bar plot of the number of differentially expressed genes (DEGs) found within the EEC cluster and partitioned by location along the crypt-villus axis. The EEC lineage shows comparable proportions of rescue and specificity between the crypt and villus compartments. (B) List of genes rescued in EECs, found in the intersection of DEGs downregulated by HFD and upregulated by VSG (corresponding with the outlined bar in (A). Despite similar degrees of rescue, EECs in the crypts versus villi show distinct genes rescued within each compartment. (C) List of genes rescued among the stem and Paneth clusters, as described in Figure 4C-D.
